# Tumor Copy Number Deconvolution Integrating Bulk and Single-Cell Sequencing Data

**DOI:** 10.1101/519892

**Authors:** Haoyun Lei, Bochuan Lyu, E. Michael Gertz, Alejandro A. Schäffer, Xulian Shi, Kui Wu, Guibo Li, Liqin Xu, Yong Hou, Michael Dean, Russell Schwartz

## Abstract

Characterizing intratumor heterogeneity (ITH) is crucial to understanding cancer development, but it is hampered by limits of available data sources. Bulk DNA sequencing is the most common technology to assess ITH, but mixes many genetically distinct cells in each sample, which must then be computationally deconvolved. Single-cell sequencing (SCS) is a promising alternative, but its limitations — e.g., high noise, difficulty scaling to large populations, technical artifacts, and large data sets — have so far made it impractical for studying cohorts of sufficient size to identify statistically robust features of tumor evolution. We have developed strategies for deconvolution and tumor phylogenetics combining limited amounts of bulk and single-cell data to gain some advantages of single-cell resolution with much lower cost, with specific focus on deconvolving genomic copy number data. We developed a mixed membership model for clonal deconvolution via non-negative matrix factorization (NMF) balancing deconvolution quality with similarity to single-cell samples via an associated efficient coordinate descent algorithm. We then improve on that algorithm by integrating deconvolution with clonal phylogeny inference, using a mixed integer linear programming (MILP) model to incorporate a minimum evolution phylogenetic tree cost in the problem objective. We demonstrate the effectiveness of these methods on semi-simulated data of known ground truth, showing improved deconvolution accuracy relative to bulk data alone.

## 1 Introduction

Cancer is one of the most lethal terminal diseases in the world, resulting for example in approximately 600,000 deaths in the U.S.A. in the past year [34]. Nevertheless, the age-adjusted rate of cancer deaths in the U.S.A. has been declining, partly due to the invention of new cancer treatments. Recent work in developing cancer therapeutics is based on the notion of personalized or precision medicine [6] to target driver alterations in specific cancer genes. Such targeted treatments have shown success in prolonging life but rarely lead to durable cures [12], largely because tumors are not normally static or homogeneous entities [8]. Most cancers exhibit phenotypes of hypermutability [20] that result in a process of continuing evolution of clonal populations of tumor cells [27], creating the opportunity for continuing acquisition of adaptive mutations as well as putatively selectively neutral genetic variants [41]. As a consequence, different cells in the same tumor may acquire distinct sets of somatic alterations, including single nucleotide variants (SNVs), copy number alterations (CNAs), and structural variations (SVs) such as gene fusions or chromosomal rearrangements. This phenomenon, called intra-tumor heterogeneity (ITH) [24], allows tumors to develop resistance to targeted treatments, as treatment-resistant subclones emerge within the tumor [27,12] or expand from initially rare subpopulations within the tumor’s clonal diversity. Considerable recent research into the molecular mechanisms of cancer has concentrated on characterizing ITH and reconstructing the processes of clonal evolution by which it develops across tumor progression (see, for example, [30]).

Currently, the most common technology to profile ITH is bulk DNA sequencing, which allows one to observe aggregate genetic variation in tumors and possibly matched normal tissue from the same patients. Bulk DNA sequencing allows one to identify reasonably common genetic lesions and estimate their variant allele fractions (VAFs). Resolving these VAFs into models of clonal heterogeneity, however, requires solving a challenging computational inference problem, known as *genomic deconvolution,* which strives to explain VAFs as mixtures of unobserved clonal sequences occurring at varying frequencies within the tumor. These methods have limited accuracy and resolution, particularly with respect to rare clonal subpopulations [1], and reveal far less clonal heterogeneity than is evident from direct single-cell analysis (e.g., [13,26]). Genomic deconvolution is particularly challenging in cancers exhibiting CNAs [38], a significant limitation given that CNAs are the primary mechanism of functional adaptation in at least some cancer types [44,21] and that CNAs at specific loci can have important consequences for treatment outcome (e.g., [25]).

Single cell sequencing (SCS) has emerged as an alternative allowing for the direct inference of clonal genotypes [26]. SCS itself is limited by difficult technical artifacts, however, such as the phenomenon of allelic dropout [14] and distortion of copy numbers due to the amplification steps used in most SCS methods to date [19]. Moreover, SCS is relatively costly in comparison to bulk sequencing. As a result, SCS studies to date have involved only small cohorts [28].

The tradeoffs between bulk sequencing and SCS have recently led to the idea that we might combine them to reconstruct ITH with both accuracy and scale [23,22], yielding improved performance in bulk data deconvolution and relative to using SCS data alone. To date, though, such work has focused on SNVs specifically. There is substantial value in developing comparable methods for CNAs given their biological importance, the greater difficulty of CNA deconvolution, and their suitability for phylogenetics from low-coverage SCS [26].

In this work, we develop methods for combining bulk and single-cell data to characterize ITH by CNAs specifically, both as a stand-alone inference and joint with phylogenetic inference on clonal subpopulations. We pose the problem of inferring the tumor subpopulations and their representation across genomic samples using a variant of non-negative matrix factorization (NMF). We seek solutions that deconvolve bulk data while achieving consistency between inferred single cells and limited SCS data. We consider two problem variants, one minimizing genomic distance between SCS-observed single-cells and inferred clones and the other explicitly incorporating a tumor phylogeny model to favor solutions that yield parsimonious evolution models relating observed cells and inferred clones. We characterize performance of the methods on semi-simulated data generated from low-coverage SCS. We show that both methods are effective at improving clonal deconvolution of CNAs with limited amounts of SCS data, with increasing accuracy as the number of genomic samples grows. We further show that explicitly modeling clonal evolution notably improves accuracy, suggesting the value of accounting for the process of tumor evolution in characterizing clonal structure.

## 2 Methods

### 2.1 Non-Negative Matrix Factorization (NMF) Deconvolution Model

As in previous work [31], we formalize the generic problem of genomic deconvolution in terms of a mixed membership model but here relating bulk and SCS data. We focus here specifically on deconvolution of copy number data, as in [38,43], which we assume is profiled on a set of *m* genomic regions. In the pure deconvolution problem, we assume a set of *n* bulk samples, which might correspond to measurements from distinct tumor sites or regions in one patient. These bulk samples are collectively encoded in an *m × n* matrix ***B***, where element *bij* corresponds to the mean copy number of locus *i* in sample *j*. Our goal is to identify an *m × k* matrix of mixture components ***C***, representing copy numbers of inferred common clones, and a ***k × n*** matrix of mixture fractions, ***F***, describing the degree to which each column of ***C*** is represented in each column of ***B. B*** is presumed to be approximated by the product of ***C*** and ***F***. We seek to minimize the deviation between ***B*** and ***C × F*** by some measure, such as the Frobenius norm. With the additional constraints that ***B***, ***C***, and ***F*** are non-negative, the problem is known as non-negative matrix factorization (NMF) [40]. More formally, we seek:

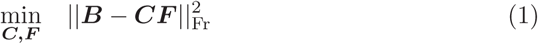

where || • ||_Fr_ is the Frobenius norm of the matrix, 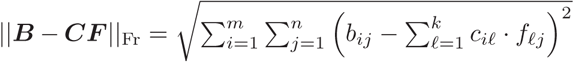, subject to the constraints 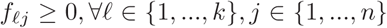; 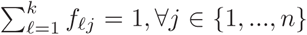; 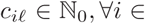 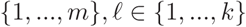.

This optimization problem is non-convex, but prior work showed that the Euclidean distance between ***B*** and ***CF*** is non-increasing under the following multiplicative update rules [18]:

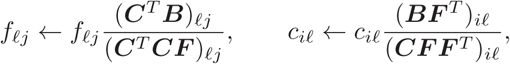

providing formulas for iterative local optimization by fixing ***C*** or ***F*** on alternate steps. In practice, we modify this process heuristically to renormalize columns of ***F*** after each iteration to ensure they add to 1. Since this heuristic might undermine the guarantee of monotonicity, we manually verify that ||***B*** — ***CF***||_Fr_ decreases on each iteration, terminating the optimization if it fails to yield continuing improvements. More details are provided in the Supplementary Methods.

Fig. 1 provides an illustrative example of the deconvolution model. Suppose we have a possible ***B***, ***C***, and ***F***. The two data points *B*_1_ and *B*_2_ represent bulk tumor samples combining three mixture components *C*_1_, *C*_2_ and *C*_3_. For ease of illustration, we assume data are assayed on the copy numbers of just two genomic loci, *G*_1_ and *G*_2_. The matrix ***B*** represents the average copy numbers of *G*_1_ and *G*_2_ in the bulk tumor samples *B*_1_ and *B*_2_. In each component of C, the copy numbers should be integers, but since the bulk tumors are weighted mixtures of components, the values in ***B*** need not be integers. The matrix ***F*** represents the fractional weights used to generate *B*_1_ and *B*_2_ from the pure components in ***C***. For example, the first column of ***F*** indicates that *B*_1_ is a mixture of equal parts of *C*_1_ and *C*_2_. This relationship can be expressed via matrix multiplication, ***B*** = ***CF***, as shown in the right part of Fig. 1.

**Fig. 1.**
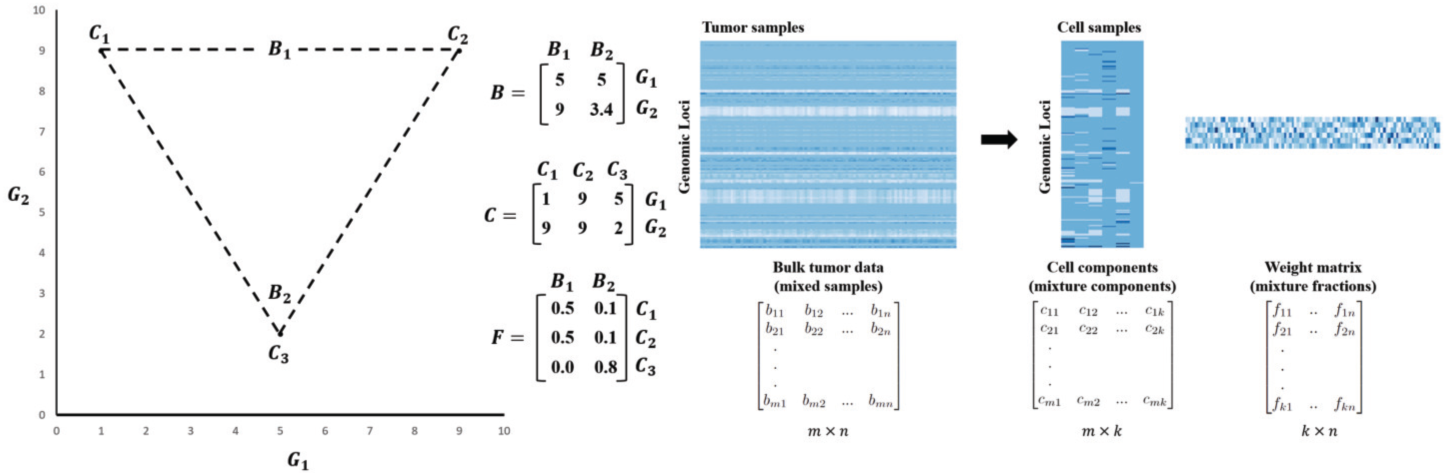
Illustration of the mathematical formulation for the mixed membership modeling problem. The model implies that each entry of ***B, C*** and ***F*** is non-negative, each entry of ***C*** is integer, and each column of ***F*** must sum up to 1.

### 2.2 Extending NMF with Single Cell Sequence (SCS) Data

The multiplicative update algorithm is a standard method for the pure NMF optimization problem, provided the number of samples n is large compared to the intrinsic dimension k of the mixture. We would, however, expect it to perform poorly for our problem, in part because real tumor data generally include few samples per patient and in part because deconvolution of copy numbers is an underdetermined problem. We sought to improve the optimization by biasing the objective function to favor inferred clones similar to the observed SCS data via an auxiliary penalty in the objective function, similar to the approach of [23,22]. Intuitively, we assume that the inferred clones (***C***) should be closely related to one or more of the observed single cells, which we call observed cell components (***C***^(*observed*)^). While any given single cell may not exactly match a consensus clone, we propose that the method will be able to approximately infer mixture components reflecting dominant clones by balancing quality of deconvolution against similarity to observed single cells. We quantify this intuition using the Euclidean distance between the inferred clones and observed cells, introducing a regularization parameter α to balance the weight of this penalty relative to the prior cost based on deconvolution quality. The resulting combined objective appears as (Eq. (2)):

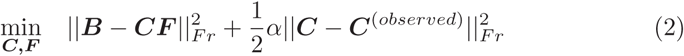

which we optimize subject to the constraints *f*_*𝓁j*_ ≥ 0, ∀𝓁 ∈ {1,…, *k*}, *j* ∈ {1,…, *n*}; 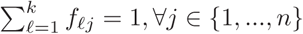; 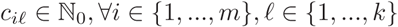.

We solve for the revised model through an extension of the iterative update algorithm [18,3]:

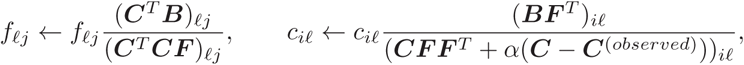

adding the constraints on ***C*** and ***F*** to the update rules [37]. We further heuristically improve on the standard practice of random initialization by initializing the cell component matrix *C* with true SCS data. Pseudocode for the complete algorithm is provided in Supplementary Methods as Algorithm 1. Collectively, these additions to the pure NMF iterative update algorithm constitute our first approach to integrating SCS data for improved deconvolution of CNAs from bulk DNA-seq, which we dub our phylogeny-free method.

### 2.3 Extending the NMF Model with a Single-Cell Phylogeny Objective

We next developed an alternative phylogeny-based approach, seeking to deconvolve the bulk data into clonal subpopulations while simultaneously inferring a phylogeny on those deconvolved clones, similar to the SNV PHiSCS method of Malikic et al. [22]. Intuitively, evolutionary distance provides a more biologically motivated measure of what we mean in asserting that inferred single cells should be similar to observed single cells. As with the phylogeny-free method, we would expect that any small sample of single cells will not exactly reflect the spectrum of dominant clones, but that the method will be able to approximately infer dominant clones by balancing deconvolution quality against evolutionary distance of mixture components to observed single cells. This approach trades off a more principled measure of solution quality for a harder optimization problem.

We quantify phylogenetic distance as the minimum over evolutionary trees incorporating both observed single cells and inferred clones of the *L*_1_ distance be-tween copy number vectors describing each tree edge. Let *C\** = [*C, C*^(*observed*)^] be a *m* × *k*\* matrix consisting of columns representing inferred clonal copy numbers followed by columns representing the copy numbers of the observed cells. Let 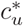 denote column *u* of *C*\*. We introduce a *k*\* × *k*\* matrix of binary variables *S*. A value of *s*_*uv*_ = 1 indicates the existence of a directed edge from node *u* to node *v*, and a value *s*_*uv*_ = 0 indicates the absence of such a edge; we set *s*_*uu*_ = 0 to avoid self loops. In other words, *S* is an adjacency matrix for a directed graph; in the full formulation (Supplementary Methods) we introduce constraints that ensure the graph is a tree. We define our measure of tree cost to be

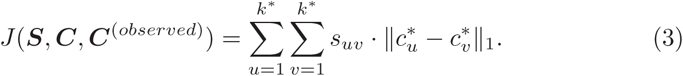

Intuitively, *J*(*S, C, C*^(*observed*)^) is a form of minimum evolution model on a phylogeny defined by *S*. While there are more sophisticated and realistic models for CNA distance (e.g.,[5,4,10]), we favored *L*_1_ distance here as a tractable approximation easily incorporated into the overall ILP framework. Similarly, while there are now a number of sophisticated methods available specifically for phylogenetics of single-cell sequences (c.f., [17]) these are largely focused on SNV rather than CNA phylogenetics (e.g.,[15,29,45]) with limited exceptions [39,36].

More specifically, we modify the NMF objective function as follows:

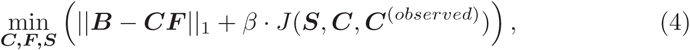

where 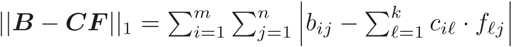 and *β* is a regularization parameter to balance deconvolution quality against parsimony of the evolutionary model. The norm ‖ • ‖_1_ is the element-wise *L*_1_ matrix norm, i.e., the sum of the absolute values of matrix elements, rather than the induced *L*_1_ matrix norm for which the same notation is sometimes used. These are optimized subject to the same constraints as in the previous formulations: 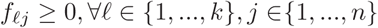; 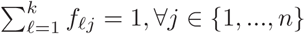; 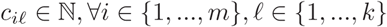.

The discrete tree optimization term lacks an analytic expression and hence does not lend itself to the prior iterative update strategy. We therefore employ a different computational strategy based on integer linear programming (ILP) to replace the linear algebra steps of the Lee and Seung method [18], similar to other recent work in joint deconvolution and phylogenetics [43,9].

For this optimization problem, we use an iterative coordinate descent approach. There are three sets of variables over which to optimize: the weight matrix *F*, the tree structure *S*, and the inferred copy numbers *C*. We solve for variables *F, S*, and *C* alternately, in this order, while holding all other variables as constant. The iterative coordinate descent continues until the decrease between successive values of *C* falls below some threshold. To initialize *C*, we used observed single cell data. Whenever two of the three sets of variables is held constant, the resulting optimization problems can each be expressed as either a linear program (LP) or an integer linear program (ILP).

When certain subsets of the variables are fixed, the resulting LP or ILP may be simplified. When solving for *F* with fixed values of *S* and *C*, the term *J*(*S, C, C*^(*observed*)^) is constant and the value of *S* is irrelevant. Similarly, when solving for *S* for fixed values of *F* and *C*, the term ‖*B* — *CF*‖_1_ is constant and therefore *F* is irrelevant. The optimal value of *C* for fixed values of *F* and *S*, however, depends both on *F* and *S*. We note that in the limit of using no single-cell data, our problem statement and method is similar to that of Zaccaria et al. [43] for incorporating tree mixtures into purely-bulk CNA deconvolution.

We solve for *S* via an ILP that uses a flow model to constrain solutions to a minimum evolution tree, adapting a similar ILP method originally developed for finding maximum parsimony character-based phylogenies [35]. Intuitively, the model forces a tree structure by setting up a flow from an arbitrary root to each other clone in the tree and minimizing the cost of edges needed to accommodate all such flows. The full ILP is described in the Supplementary Methods.

### 2.4 Validation via Observed Single-Cell Data

To validate the method, we require bulk data for which clone copy number vectors and frequencies are known. As this is unavailable for any real dataset, we use semi-simulated data generated from CNV calls [2] from real SCS data from two human glioblastoma cases [42]. The full single cell data set consists of low-depth SCS DNA-seq used to establish mean copy numbers at 9934 genomic positions throughout the genome, at intervals of approximately 40kbp. Each tumor was subdivided into three regions (i.e., samples), with each single cell labeled by its region (1, 2, or 3) of origin. We used these true SCS CNA data to generate a series of synthetic bulk data sets, simulating either one, two, or three bulk samples from each region for a total of three, six, or nine bulk samples per trial. Each simulated sample is generated by sampling two dominant cells from a region to represent major clones, twenty three other cells from the same region to represent minor clones, and 50 cells from the other regions to represent contamination, which are mixed with Dirichlet-sampled proportions with weight parameters for dominant, minor, and contaminant clones in the ratio 10 to 0.1 to 0.01. We then assessed our ability to deconvolve the bulk data across a range of regularization parameter values and random replicates of the chosen single cells. We assessed accuracy by the fraction of genomic positions assigned correct copy number and by the root mean square deviation (RMSD) between true and inferred cell components and mixture fractions. Fig. 2 summarizes the overall experimental design, which is described in more detail in the Supplementary Methods in Sec. A.4. This design treats observed SCS as the ground truth, allowing us to ignore the problem of doublet cells that typically must be addressed with SCS data. We would normally require that likely doublets be removed from SCS data in preprocessing before applying our method. This design also does not explicitly include calling CNA markers on bulk data, itself a hard problem that would need to be performed in preprocessing before applying our method.

**Fig. 2.**
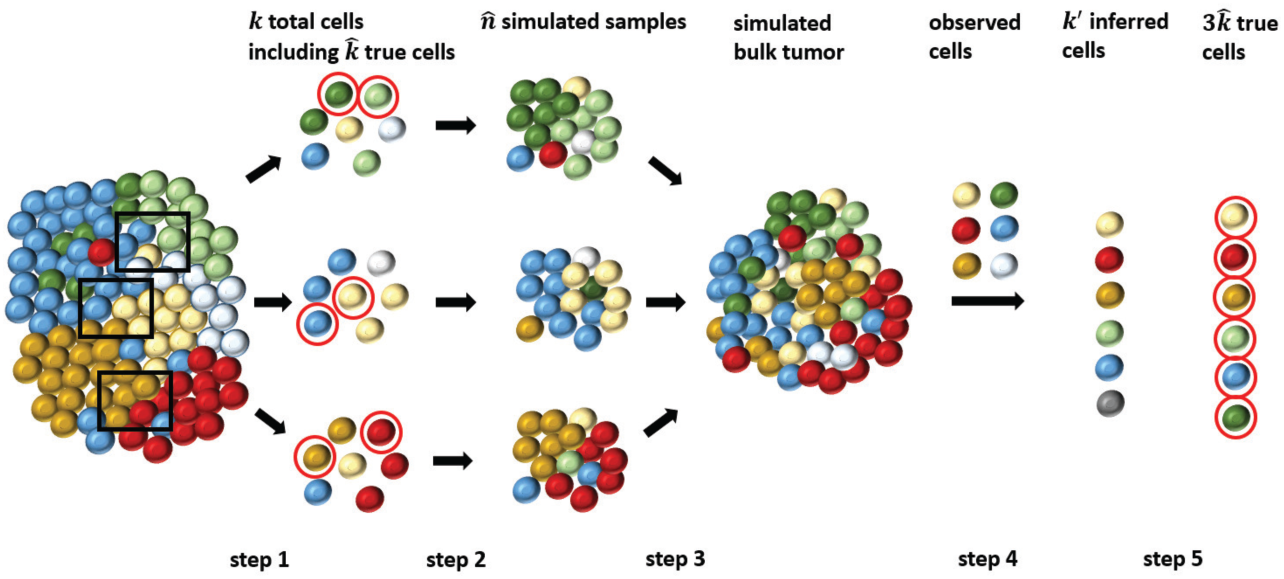
Work-flow for the simulation and validation. We separate the whole process into 5 main steps: in step 1, we randomly chose *k* total single cells from each region (indicated by the black frames), where we can pick 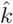 dominant clones (indicated by red circles, also called true cells); in step 2, we simulated 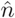 tumor samples from each region using the *k* cells; in step 3, we combined the 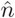 tumor samples to get a simulated bulk tumor; in step 4, we deconvolved the bulk tumor integrating observed cells to get 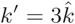 inferred clones; and in step 5, we assessed the performance using the *k*′ inferred clones and 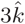 true cells.

We were unable to identify any competitive tool for bulk deconvolution of purely CNA data applicable to small numbers of bulk samples and for which software is publicly available. We therefore compare our methods to standard NMF, as implemented by our code with zero regularization parameters.

### 2.5 Implementation

The methods described in Methods and refined below were all implemented in Python3, using Gurobi. One practical change from the formulation above is that we replaced the theoretical *f*_*𝓁j*_ ≥ 0 with *f*_*𝓁j*_ > 10^-4^ to avoid having the *f* values trapped at 0. The observed human subjects data cannot be re-distributed, but code for the methods is available along with artificial data on Github (https://github.com/CMUSchwartzLab/SCS_deconvolution).

## 3 Results

### 3.1 Phylogeny-Free Method

We first assessed the accuracy of the phylogeny-free method relative to pure NMF and simple heuristic improvements. Fig. 3 provides a summary of accuracy and RMSD for inference of true SCS components via the method of Sec. 2.2 for variations in the number of tumor samples (3, 6, 9) and regularization parameter *α* (0-1) over 40 replicates per condition. To provide a baseline for comparison, each plot provides equivalent accuracy measures for NMF [18] (i.e., Algorithm 1 with *α* = 0) with random initial integer valued *C* (red dashed line in Fig. 3) and with the proposed solution that all copy numbers have the normal value of 2, which we call the “all-diploid baseline” (black dashed line in Fig. 3 and Fig. S3). In each case, the bulk data is simulated from *k*′ = 6 fundamental cell components (2 out of a random 25 cells selected in each region).

**Fig. 3.**
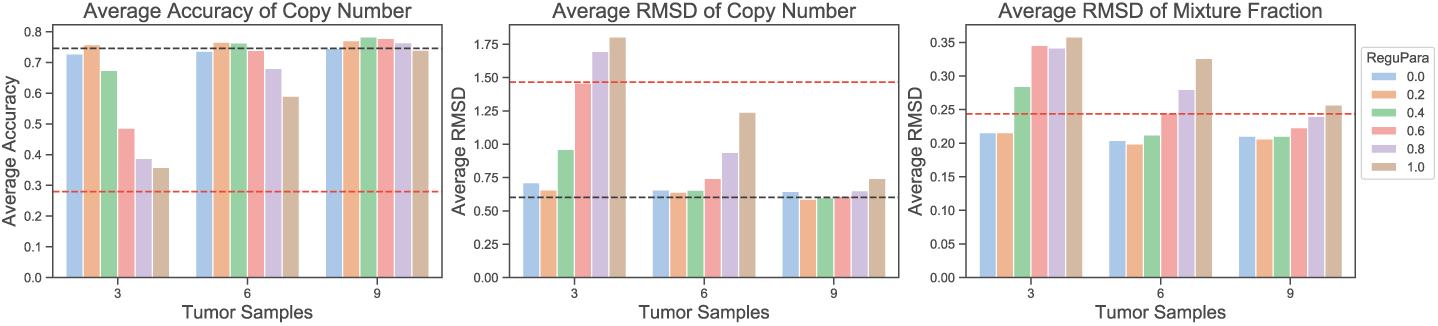
Accuracy and RMSD of the phylogeny-free method as functions of tumor sam-ples and regularization parameter. The red dashed line shows average overall accuracy (left panel) or RMSD (center and right panels) of NMF with random initialization. The black dashed line shows the performance of the all-diploid baseline solution. Since we cannot resolve mixture fractions for an all-diploid solution, we omit it from the mixture fraction results. Different bars show performance as a function of regulariza-tion parameter *α* of Eq. 2 from 0.0 to 1.0 in increments of 0.2. The X-axis shows the number of tumor samples and the Y-axis the average accuracy or RMSD.

Pure NMF with random initialization performed poorly, which is unsurprising since NMF on CNA data is an underdetermined problem, although the simple heuristic of biasing the search toward biologically plausible solutions by initializing with real SCS data improves accuracy. Bringing true SCS data into the objective function yielded modest improvements in accuracy over using SCS data solely for initialization for at least some values of the regularization parameter. The phylogeny-free method with *α* = 0 corresponds to pure NMF initialized with true SCS data, and this performed slightly worse than the all-diploid baseline solution. Modestly increasing *α* led to some improvement in accuracy, but above some value, *α* put too much weight on similarity to observed SCS data and too little weight on quality of the deconvolution, giving worse overall results. The best value of *α* depended on sample size, which we attribute again to NMF being underdetermined if the number of desired components is larger than the number of samples. The plots suggest that the method is fairly robust to *α* if the number of samples exceeds the intrinsic dimension of the data (six), but that SCS data can overcome that limit for small numbers of samples with a well-tuned regularization term. Additional Supplementary Results show minimal additional improvement even with unrealistically large sample sizes (Fig. S4), and also show the performance is consistent across individual inferred clones (Fig. S5).

Fig. 4 provides an illustrative example of performance for a single selected clone inferred from three, six, or nine samples, intended to demonstrate kinds of errors the method tends to produce. We chose the one cell component with smallest RMSD for each sample size to simplify visual inspection. We see that at least in these high-quality cases, the distributions of copy numbers are similar for the inferred and true cells. For loci at or just above diploid, the modified NMF can usually infer the exact copy number. Where errors occur, they tend to be in loci with large (5-10) or smaller copy numbers (0-1).

**Fig. 4.**
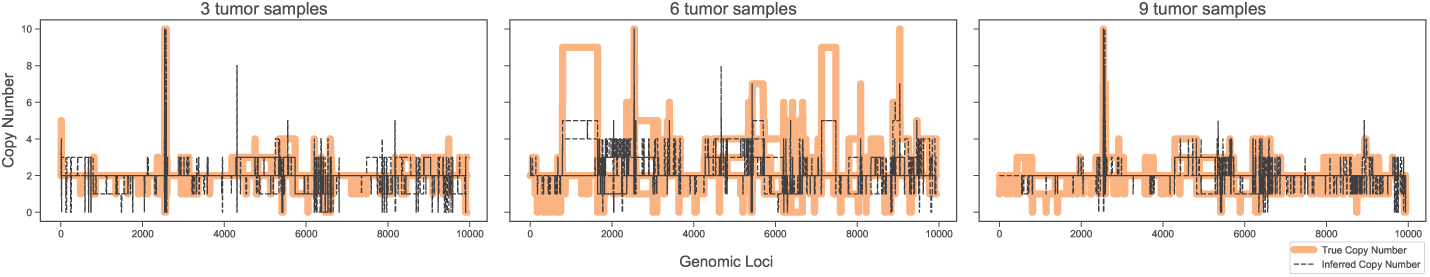
Visualization of copy number as a function of genomic locus for single examples of inferred and true clones for the phylogeny-free method for three, six, and nine samples. The figure uses the minimum-RMSD pair for each case. The black dashed line shows the copy number inferred by modified NMF and the orange bar shows the true copy number in that position.

### 3.2 Phylogeny-based Method

We next examined results of the phylogeny-based method of Sec. 2.3 under the same conditions used to assess the phylogeny-free method. Fig. 5 summarizes average accuracy and RMSD as a function of regularization parameter *β*. The figure compares the results of pure NMF with the all-diploid baseline. Setting *β* = 0 provides poor performance, substantially below the all-diploid baseline solution. Making *β* = 0 for Fig. 5 represents the same optimization problem as *α* = 0 for Fig. 3, but solved by the coordinate descent method we developed to accommodate the ILP phylogeny objective rather than by the modified iterative update algorithm with the simpler *L*_2_ objective. Fig. 5 thus suggests that the new coordinate descent method is less effective at pure NMF than is the prior iterative update algorithm. Despite that observation, the results on *β* ≥ 0.2 show substantially better accuracy than was achieved by pure NMF or the phylogeny-free algorithm. Further, the results appear robust to variation in *β* across the range examined. Supplementary Results distinguishing accuracy across cells (Fig. S6) support the robustness of the phylogeny-based method to a range of *β* values in cell-to-cell inferences.

**Fig. 5.**
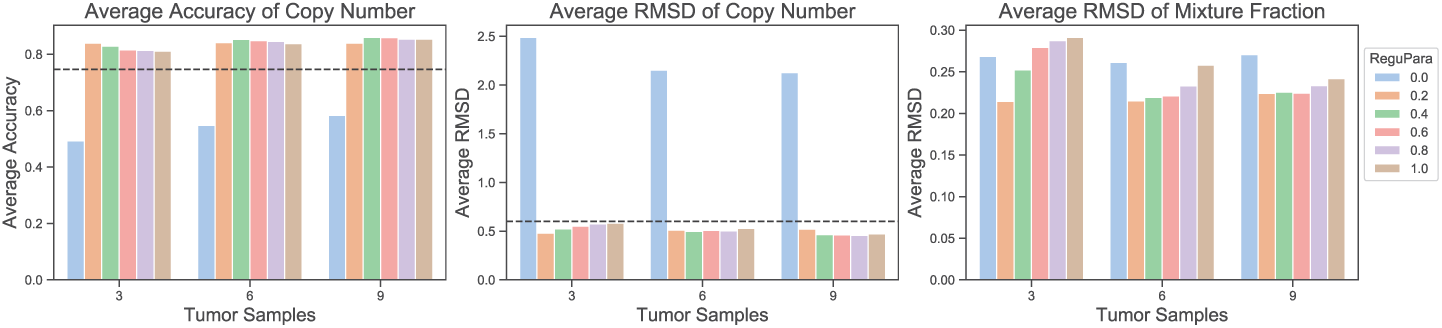
Average accuracy and RMSD for the phylogeny-based method as functions of tumor samples and regularization parameter. The left panel shows the average accuracy of inferred copy numbers, the center panel average RMSD between inferred and true copy numbers, and the right panel average RMSD between the inferred and true mixture fractions. The black dashed line shows the performance of the all-diploid baseline solution. Since we cannot resolve mixture fractions for an all-diploid baseline, we omit it from the mixture fraction results. Bar plots show performance with different regularization parameters β of Eq. 3 from 0.0 to 1.0 with increment of 0.2. The X-axis shows the number of tumor samples and the Y-axis the average accuracy or RMSD.

Fig. 6 shows copy numbers for a single minimum-RMSD pair for inferred and true clones for each number of samples, again to visualize the nature of inference errors. The results again show exact fitting for most loci, as well as better fitting for both large (5-10) and small (0-1) copy numbers than the phylogeny-free method of Fig. 4. There is no evident pattern to the smaller number of errors that do occur for the phylogeny-based versus phylogeny-free method, which are observed for a range of low and high copy number values.

**Fig. 6.**
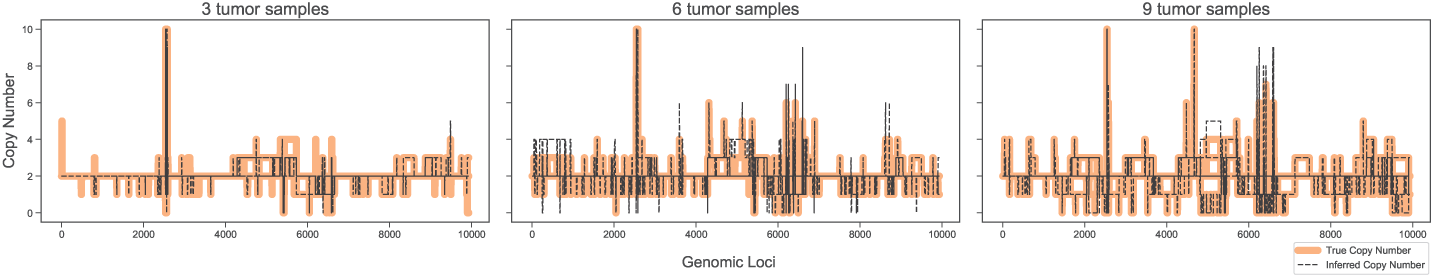
Visualization of copy number as a function of genomic locus for single examples of inferred and true clones for the phylogeny-based method for three, six, and nine samples. The figure uses the minimum-RMSD pair for each case. The black dashed line shows the inferred copy number and the orange bars show the true copy number in each position.

Fig. 7 compares the two methods at their optimal regularization parameters for three, six, and nine tumor samples. The phylogeny-based method outperforms the phylogeny-free method in accuracy and copy number RMSD in all cases. It is slightly better in mixture fraction RMSD for three samples, but worse for six and nine samples. Fig. S7 in the Supplementary Results shows comparative performance of the two methods in individual cell components. Given the poorer performance at pure NMF of the phylogeny-based method’s algorithm versus the phylogeny-free method’s algorithm, we tentatively attribute the phylogeny-based method’s better overall performance to better evolutionary distance estimates and not to a better optimization algorithm.

**Fig. 7.**
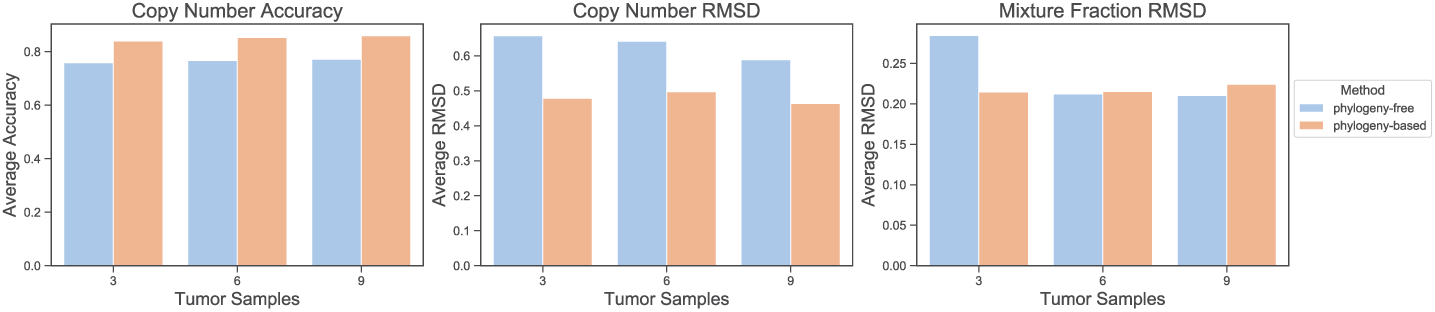
Comparison between phylogeny-free and phylogeny-based methods. Bar graphs show average accuracy and RMSD over all cell components and replicates using the optimal regularization parameter for the given method, measure, and number of samples. The left panel shows accuracy in copy numbers for α=0.2, 0.2, 0.4 for the phylogeny-free method and, *β*=0.2, 0.4, 0.4 for the phylogeny-based method for 3, 6 and 9 tumor samples, respectively. The center panel shows RMSD of copy numbers for α=0.2, 0.2, 0.2 for the phylogeny-free method and β=0.2, 0.4, 0.4 for the phylogeny-based method for 3, 6, and 9 tumor samples, respectively. The right panel shows RMSD of mixture fractions for α=0.2, 0.2, 0.2 for the phylogeny-free method and β=0.2, 0.2, 0.2 for the phylogeny-based method for 3, 6, and 9 tumor samples, respectively. The X-axis shows the number of tumor samples and the Y-axis the average accuracy or RMSD.

The phylogeny-based method also provides as output the phylogeny. While we cannot exhaustively show trees across all replicates, we provide three representative examples in Fig. 8. Because we use true SCS data to generate our synthetic mixtures, we do not know the full ground truth trees for the data and do not attach any biological meaning to the inferred trees. We can partially validate correctness of the trees using the fact that the cells were gathered from distinct tumor regions, and while we would not expect clonal ancestry to segregate perfectly by region we should see a trend towards closer evolutionary relationship among cells in spatial proximity. We tested whether pairs of cells from distinct regions cluster together in disjoint subtrees (a kind of partial-information quartet distance); we found that a significant majority of pairs-of-pairs do (79% for 3-sample data, 74% each for 6- or 9-sample data) providing some support for the biological relevance of the trees.

**Fig. 8.**
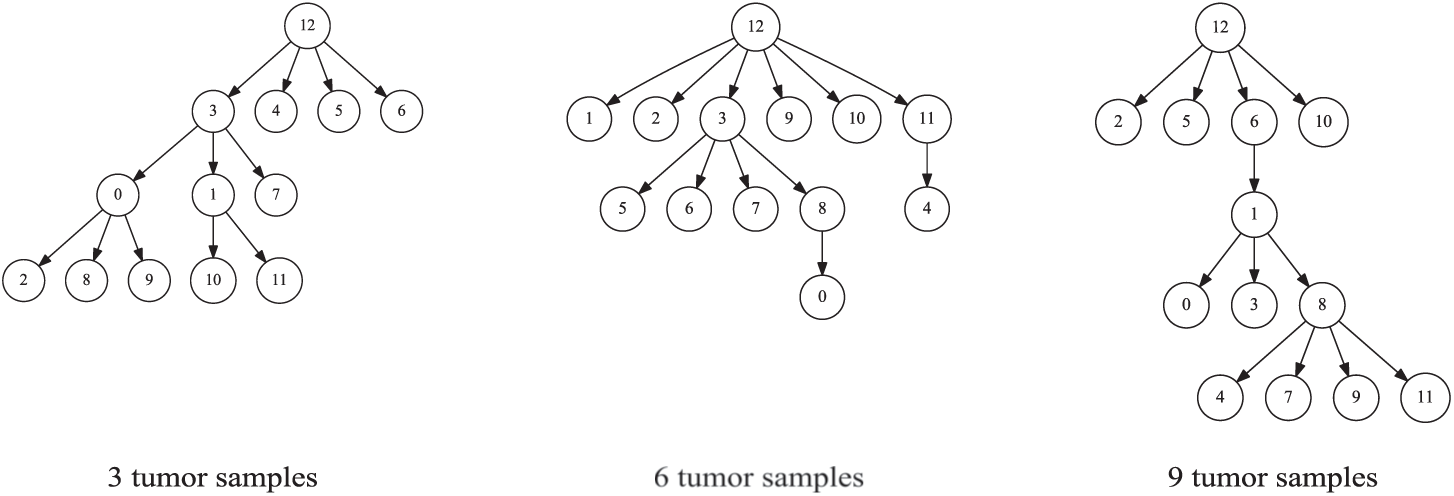
Tree structure inferred via the phylogeny-based method ILP method for three problem instances. The examples come from the same instances used to pick the representative copy number profiles in Fig. 6. In each tree, nodes 0-5 are inferred cells, nodes 6-11 are observed cells, and node 12 is the diploid root.

## 4 Conclusions and Discussion

We presented two novel methods for deconvolving clonal copy number variation from bulk tumor genomic data assisted by small amounts of SCS data. The work is intended to provide a practical strategy for producing high-quality clonal CNA deconvolution scalable to large tumor cohorts in the face of still high costs of single-cell DNA sequencing. Validation on semi-simulated data shows that limited amounts of SCS copy number data can be productively used to improve upon pure bulk deconvolution, as assessed by accuracy in inferring clonal copy number profiles and their proportions in single- or multi-sample tumor genomic data. We showed substantial improvement by explicitly constructing clonal phylogenies jointly with deconvolution, suggesting the value of a principled evolutionary model in inferring accurate clonal structure.

While this work provides a proof-of-principle demonstration for combining bulk and SCS data for CNA deconvolution, it also suggests a need for future work. Data of the kind needed by this study remain rare, largely because current SCS studies have not been designed for such a hybrid approach. Most studies to date have profiled many single cells from few patients rather than few cells from larger cohorts, as the current work proposes. We hope that demonstrating the effectiveness of the strategy will promote its use in future study designs, and stimulate new thinking on how most effectively to use single-cell sequencing technologies to solve the underlying data science problems, in turn creating more data on which similar algorithms can be improved. The framework might also be improved in a variety of ways, including more realistic tree models and consideration of other constraints one can extract from SCS data. For example, we considered only penalty terms on ***C*** but might also use SCS to improve estimates of the clonal frequency matrix ***F*** [33]. The method might also be improved by replacing L1 distance with measures reflecting more sophisticated models of CNA-driven evolution [5,32,4,10]. It could be useful to identify minor clones that have likely loss of heterozygosity events, since these may influence clinical outcomes, and to automate inference of the number of dominant clones. It would also be useful to combine the CNAs of this work with the SNVs of Malikic et al. [23,22], as is commonly done now for bulk deconvolution (e.g., [7,16,11]), and to leverage more effectively data from new low-coverage SCS DNA-seq methods [46] or long-read sequencing. In addition, our algorithms for solving for these models are heuristic and we might productively consider alternative methods to approach true global optima or to improve scalability to larger datasets.

## Acknowledgements

This research was supported in part by the Intramural Research Program of the National Institutes of Health, National Library of Medicine and both Center for Cancer Research and Division of Cancer Epidemiology and Genetics within the National Cancer Institute. This research was supported in part by the Exploration Program of the Shenzhen Science and Technology Innovation Committee [JCYJ20170303151334808]. Portions of this work have been funded by U.S. N.I.H. award R21CA216452 and Pennsylvania Dept. of Health award 4100070287. The Pennsylvania Department of Health specifically disclaims responsibility for any analyses, interpretations or conclusions.

## References

1. Barber, L.J., Davies, M.N., Gerlinger, M.: Dissecting cancer evolution at the macroheterogeneity and micro-heterogeneity scale. Current Opinion in Genetics and Development 30, 1–6 (2015)

2. Baslan, T., Kendall, J., Rodgers, L., Cox, H., Riggs, M., Stepansky, A., Troge, J., Ravi, K., Esposito, D., Lakshmi, B., et al.: Genome-wide copy number analysis of single cells. Nature Protocols 7(6), 1024 (2012)

3. Berry, M.W., Browne, M., Langville, A.N., Pauca, V.P., Plemmons, R.: Algorithms and applications for approximate nonnegative matrix factorization. Computational Statistics & Data Analysis 52(1), 155–173 (2007)

4. Chowdhury, S.A., Gertz, E.M., Wangsa, D., Heselmeyer-Haddad, K., Ried, T., Schaffer, A., Schwartz, R.: Inferring models of multiscale copy number evolution for single-tumor phylogenetics. Bioinformatics 31(12), i258–i267 (2015)

5. Chowdhury, S., Shackney, S., Heselmeyer-Haddad, K., Ried, T., Schaffer, A., Schwartz, R.: Algorithms to model single gene, single chromosome, and whole genome copy number changes jointly in tumor phylogenetics. PLoS Computational Biology 10(7), e1003740 (2014)

6. Coyne, G.O., Takebe, N., Chen, A.P.: Defining precision: The precision medicine initiative trials NCI-IMPACT and NCI-MATCH. Current Problems in Cancer 41, 182–193 (2017)

7. Deshwar, A.G., Vembu, S., Yung, C.K., Yang, G.H., Stein, L., Morris, Q.: Phy-loWGS: Reconstructing subclonal composition and evolution from whole-genome sequencing of tumors. Genome Biology 16, 35 (2015)

8. Dexter, D.L., Leith, J.T.: Tumor heterogeneity and drug resistance. Journal of Clinical Oncology 4(2), 244–257 (1986)

9. Eaton, J., Wang, J., Schwartz, R.: Deconvolution and phylogeny inference of structural variations in tumor genomic samples. Bioinformatics 34, i357–i365 (2018)

10. El-Kebir, M., Raphael, B., Shamir, R., Sharan, R., Zaccaria, S., Zehavi, M., Zeira, R.: Complexity and algorithms for copy-number evolution problems. Algorithms for Molecular Biology 12(1), 13 (2017)

11. El-Kebir, M., Satas, G., Oesper, L., Raphael, B.J.: Inferring the mutational history of a tumor using multi-state perfect phylogeny mixtures. Cell Systems 3(1), 43–53 (2016)

12. Fisher, R., Pusztai, L., Swanton, C.: Cancer heterogeneity: implications for targeted therapeutics. British Journal of Cancer 108(3), 479–485 (2013)

13. Heselmeyer-Haddad, K., Berroa Garcia, L.Y., Bradley, A., Ortiz-Melendez, C., Lee, W., Christensen, R., Prindiville, S.A., Calzone, K.A., Soballe, P.W., Hu, Y., et al.: Single-cell genetic analysis of ductal carcinoma in situ and invasive breast cancer reveals enormous tumor heterogeneity yet conserved genomic imbalances and gain of MYC during progression. American Journal of Pathology 181(5), 1807–1822 (2012)

14. Hou, Y., Song, L., Zhu, P., Zhang, B., Tao, Y., Xu, X., Li, F., Wu, K., Liang, J., Shao, D., et al.: Single-cell exome sequencing and monoclonal evolution of a JAK-2 negative myeloproliferative neoplasm. Cell 148(5), 873–885 (2012)

15. Jahn, K., Kuipers, J., Beerenwinkel, N.: Tree inference for single-cell data. Genome Biology 17(1), 86 (2016)

16. Jiang, Y., Qiu, Y., Minn, A.J., Zhang, N.R.: Assessing intratumor heterogeneity and tracking longitudinal and spatial clonal evolutionary history by next-generation sequencing. Proceedings of the National Academy of Sciences 113(37), E5528-E5537 (2016)

17. Kuipers, J., Jahn, K., Beerenwinkel, N.: Advances in understanding tumour evolution through single-cell sequencing. Biochimica et Biophysica Acta (BBA)-Reviews on Cancer 1867(2), 127–138 (2017)

18. Lee, D.D., Seung, H.S.: Algorithms for non-negative matrix factorization. In: Advances in Neural Information Processing Systems. pp. 556–562 (2001)

19. Lei, H., Ma, F., Chapman, A., Lu, S., Xie, X.S.: Single-cell whole-genome amplification and sequencing: Methodology and applications. Annual Review of Genomics and Human Genetics 16, 79–102 (2015)

20. Loeb, L.A.: A mutator phenotype in cancer. Cancer Research 61(8), 3230–3239 (2001)

21. Macintyre, G., Goranova, T.E., De Silva, D., Ennis, D., Piskorz, A., Eldridge, M., Sie, D., Lewsley, L., Hanif, A., Wilson, C., et al.: Copy number signatures and mutational processes in ovarian carcinoma. Nature Genetics 50(9), 1262–1270 (2018)

22. Malikic, S., Ciccolella, S., Mehrabadi, F.R., Ricketts, C., Rahman, M.K., Haghshenas, E., Seidman, D., Hach, F., Hajirasouliha, I., et al.: PhlSCS-a combinatorial approach for sub-perfect tumor phylogeny reconstruction via integrative use of single cell and bulk sequencing data. bioRxiv p. 376996 (2018)

23. Malikic, S., Jahn, K., Kuipers, J., Sahinalp, C., Beerenwinkel, N.: Integrative inference of subclonal tumour evolution from single-cell and bulk sequencing data. bioRxiv p. 234914 (2017)

24. Marusyk, A., Polyak, K.: Tumor heterogeneity: causes and consequences. Biochimica et Biophysica Acta (BBA)-Reviews on Cancer 1805(1), 105–117 (2010)

25. McGranahan, N., Rosenthal, R., Hilley, C.T., Rowan, A.J., Watkins, T.B.K., Wilson, G.A., Birkbak, N.J., Veeriah, S., Van Loo, P., Herrero, J., et al.: Allele-specific HLA loss and immune escape in lung cancer evolution. Cell 171(6), 1259–1271 (2017)

26. Navin, N., Kendall, J., Troge, J., Andrews, P., Rodgers, L., McIndoo, J., Cook, K., Stepansky, A., Levy, D., Esposito, D., et al.: Tumour evolution inferred by single-cell sequencing. Nature 472(7341), 90–94 (2011)

27. Nowell, P.C.: The clonal evolution of tumor cell populations. Science 194(4260), 23–28 (1976)

28. Ortega, M.A., Poirion, O., Zhu, X., Huang, S., Wolgruber, T.K., Sebra, R., Garnire, L.X.: Using single-cell multiple omics approaches to resolve tumor heterogeneity. Clinical and Translational Medicine 6, 46 (2017)

29. Ross, E.M., Markowetz, F.: OncoNEM: inferring tumor evolution from single-cell sequencing data. Genome Biology 17(1), 69 (2016)

30. Schwartz, R., Schaffer, A.A.: The evolution of tumour phylogenetics: principles and practice. Nature Reviews Genetics 18(4), 213–229 (2017)

31. Schwartz, R., Shackney, S.E.: Applying unmixing to gene expression data for tumor phylogeny inference. BMC Bioinformatics 11(1), 42 (2010)

32. Schwarz, R.F., Ng, C.K., Cooke, S.L., Newman, S., Temple, J., Piskorz, A.M., Gale, D., Sayal, K., Murtaza, M., Baldwin, P.J., et al.: Spatial and temporal heterogeneity in high-grade serous ovarian cancer: a phylogenetic analysis. PLoS Medicine 12(2), e1001789 (2015)

33. Shackleton, M., Quintana, E., Fearon, E.R., Morrison, S.J.: Heterogeneity in cancer: cancer stem cells versus clonal evolution. Cell 138(5), 822–829 (2009)

34. Siegel, R.L., Miller, K.D., Fedewa, S.A., Ahnen, D.J., Meester, R., Barzi, A., Jemal, A.: Colorectal cancer statistics, 2017. CA: A Cancer Journal for Clinicians 67(3), 177–193 (2017)

35. Sridhar, S., Lam, F., Blelloch, G.E., Ravi, R., Schwartz, R.: Efficiently finding the most parsimonious phylogenetic tree via linear programming. In: International Symposium on Bioinformatics Research and Applications. pp. 37-48. Springer (2007)

36. Subramanian, A., Schwartz, R.: Reference-free inference of tumor phylogenies from single-cell sequencing data. BMC Genomics 16(11), S7 (2015)

37. Thurau, C., Kersting, K., Bauckhage, C.: Convex non-negative matrix factorization in the wild. In: 2009 Ninth IEEE International Conference on Data Mining. pp. 523–532 (Dec 2009). https://doi.org/10.1109/ICDM.2009.55

38. Tolliver, D., Tsourakakis, C., Subramanian, A., Shackney, S., Schwartz, R.: Robust unmixing of tumor states in array comparative genomic hybridization data. Bioinformatics 26(12), i106–i114 (2010)

39. Wang, Y., Waters, J., Leung, M.L., Unruh, A., Roh, W., Shi, X., Chen, K., Scheet, P., Vattathil, S., Liang, H., et al.: Clonal evolution in breast cancer revealed by single nucleus genome sequencing. Nature 512(7513), 155–160 (2014)

40. Wang, Y., Zhang, Y.: Nonnegative matrix factorization: A comprehensive review. IEEE Transactions on Knowledge and Data Engineering 25(6), 1336–1353 (2013)

41. Williams, M.J., Werner, B., Barnes, C.P., Graham, T.A., Sottoriva, A.: Identification of neutral tumor evolution across cancer types. Nature Genetics 48(3), 238–244 (2016)

42. Wu, K., Wan, W., Hou, Y., Li, F., Li, G., Xu, L., Shi, X., Dean, M., Zhang, L.: Diverse evolutionary dynamics in glioblastoma inference by multi-region and singlecell sequencing. Journal of Clinical Oncology 34(15_suppl), 11580–11580 (2016)

43. Zaccaria, S., El-Kebir, M., Klau, G.W., Raphael, B.J.: Phylogenetic copy-number factorization of multiple tumor samples. Journal of Computational Biology 25(7), 689–708 (2018)

44. Zack, T.I., Schumacher, S.E., Carter, S.L., Cherniack, A.D., Saiksena, G., Tabak, B., Lawrence, M.S., Zhang, C.Z., Wala, J., Mermel, C.H., et al.: Pan-cancer patterns of somatic copy number alteration. Nature Genetics 45(10), 1134–1140 (2013)

45. Zafar, H., Tzen, A., Navin, N., Chen, K., Nakhleh, L.: SiFit: inferring tumor trees from single-cell sequencing data under finite-sites models. Genome Biology 18(1), 178 (2017)

46. Zahn, H., Steif, A., Laks, E., Eirew, P., VanInsberghe, M., Shah, S., Aparicio, S., Hansen, C.: Scalable whole-genome single-cell library preparation without preamplification. Nature Methods 14(2), 167 (2017)

