## Supplementary Materials for "Tumor Copy Number Deconvolution Integrating Bulk and Single-Cell Sequencing Data"

### A Supplementary Methods

#### A.1 Phylogeny-free Method for Integrating SCS Data into NMF

We solve for the phylogeny-free variant of SCS-assisted NMF (Eq. 2) via an implementation of the iterative update algorithm, for which we present pseudocode as Algorithm 1. The core of the algorithm consists of the modified update rules from Sec. 2.2. Additional heuristic modifications are used to enforce non-negativity and integrality of solutions and make use of SCS data to bias initialization towards biologically plausible solutions, as well as to better handle possible numerical errors arising from finite machine precision.

**Algorithm 1:** Modified Multiplicative Update Algorithm for NMF

$\boldsymbol{C_0}$ = real single cell;
$\boldsymbol{F_0}$ = rand(k, n);
normalize $\boldsymbol{F_0}$ to make each column sum up to 1;
distance = $+\infty$;
$i = 0$;
dnorm = 1;
$\text{dnorm}_0 = 0$;
**while** distance > threshold **do**
  dnorm = $||\boldsymbol{B} - \boldsymbol{CF}||^2_{Fr}$;
  **if** $\text{dnorm}_0 - \text{dnorm} > 0$ *or* $i >$ Maxiter **then**
    quit the loop
  **end**
  numerator = $\boldsymbol{C_0^T B}$;
  $\boldsymbol{F} = \max(0, \boldsymbol{F_0} .* (\text{numerator} ./ (\boldsymbol{C_0^T C_0 F_0} + 10^{-9})))$;
  normalize $\boldsymbol{F}$ to make each column sum up to 1;
  numerator = $\boldsymbol{BF}^T$;
  $\boldsymbol{C} =$
   $\max(0, \boldsymbol{C_0} .* (\text{numerator} ./ (\boldsymbol{C_0 F F}^T + \alpha(\boldsymbol{C_0} - \boldsymbol{C}^{(observed)}) + 10^{-9})))$;
  round entries of $\boldsymbol{C}$ to the nearest integers;
  $\boldsymbol{C_0} = \boldsymbol{C}$;
  $\boldsymbol{F_0} = \boldsymbol{F}$;
  $\text{dnorm}_0 = ||\boldsymbol{B} - \boldsymbol{C_0 F_0}||^2_{Fr}$;
  distance = $\text{dnorm}_0$;
  $i \leftarrow i + 1$;
**end**

#### A.2 ILP for Phylogeny Inference in the Phylogeny-Based Method

The major change in the phylogeny-based method of Sec. 2.3 is the introduction of a minimum-evolution phylogeny cost, $J(\boldsymbol{S}, \boldsymbol{C}, \boldsymbol{C}^{(observed)})$, to the objective function (Eq. 4). We solve for the phylogeny on each pass of the coordinate descent algorithm via a multi-commodity flow ILP [35]. Let the vertex set $T = \{1, \ldots, k^*\}$ represents the set of all cells in $\boldsymbol{C^*}$. Let $r$ be the unique, predetermined, root of $T$, which in the present practice is a purely diploid node. Further, let $w_{u,v}$ be the $L_1$ distance between the copy number vectors corresponding to nodes $u, v \in T$. For $t, u, v \in T$, introduce the binary variables $g^t_{v,u}$ representing the amount of flow along edge $(u, v)$ with destination $t \in T$. The full ILP is then as follows:

$$\min \sum_{u,v} s_{uv} w_{uv} \text{ s.t.} \tag{5}$$

$$\begin{aligned}
\sum_v g^t_{uv} &= \sum_v g^t_{vu}, \ \forall u \in T, u \neq t, u \neq r \\
\sum_v g^t_{vt} &= 1, \ \forall t \in T, t \neq r \\
g^t_{vr} &= 0, \ \forall v \\
\sum_v g^t_{tv} &= 0, \sum_v g^t_{rv} = 1, \ \forall t \in T \\
0 \leq g^t_{uv} &\leq s_{uv}, \ \forall t \in T \\
s_{uu} &= 0, \ \forall u \\
s_{uv} &\in \{0, 1\}
\end{aligned}$$

Intuitively, the method defines a flow from a single root to every other vertex and requires the graph $T$ to contain edges $(u, v)$, indicated by $s_{uv} = 1$, such that all such flows can be accommodated. This forces the graph to be connected. We can further establish that the resulting graph is acyclic. For purposes of contradiction, assume the optimal $T$ contains a cycle. We can then remove any edge $(u, v)$ on the cycle and reroute any flow using that edge through the cycle in the other direction, reducing the cost of the tree by $w_{uv}$. Since $w_{uv}$ is a non-negative $L_1$ distance, then this must reduce the cost of the tree provided $u \neq v$, showing that $T$ was non-optimal. This establishes by contradiction that the optimal $T$ is connected and acyclic, i.e., a tree. It is specifically a tree of minimum cost over the complete graph of observed single cells and inferred clones implied by $\boldsymbol{C}$ and $\boldsymbol{C}^{(observed)}$.

At any stage of the algorithm, if there are multiple minimum-cost solutions for either $\boldsymbol{F}$, $\boldsymbol{S}$ or $\boldsymbol{C}$, any solution might be chosen.

#### A.3 Generation of Single-Cell Sequence (SCS) Glioblastoma Data

Data for this analysis are provided from a study of single-cell genomics in two glioblastoma patients [42]. Each patient's primary tumor was divided into three tumor regions, with 59-82 single cells extracted from each region for sequencing, for a total of 448 cells. Nuclei of 432 of these cells were amplified by multiple displacement amplification (MDA) and sequenced to a coverage of 0.17X. We used 393 among the 410 cells that passed quality control. These cells were called for CNVs by modified variable binning [2]. The resulting calls provided the input for single cell analysis and for construction of semi-synthetic bulk data in the present work.

#### A.4 Generating Semi-synthetic Data from SCS Samples

This section provides additional detail on the generation of semi-synthetic data from SCS samples for use in validating the methods. For each simulated tumor, we generate a set of experiments in which we simulated either one, two or three bulk samples from each of three tumor regions, for a total of three, six or nine bulk samples, respectively. To generate the simulated bulk samples, we randomly chose 25 single cells from each region, for a total of 75 cells. These 75 cell define the copy number vectors that make a nonzero contribution to any of the simulated bulk samples. For each region, we chose two cells from among the 25 to represent co-dominant clones. We refer to the two chosen cells as *dominant cells*. To model the noisy nature of bulk tumor data, the two dominant cells in the region, the 23 remaining cells from the same region, and the 50 cells from the other two regions all contribute to the copy number of each bulk sample, but at different mixture fractions. The two dominant cells from each region make by far the greatest expected contribution to the simulated bulk samples for that region. This design is intended to approximate clonal structure of real tumor samples, where one might see a small number of dominant clones and a long tail of rarer cell populations [13]. We assess the methods on their ability to infer the dominant clones, with the rare clones effectively serving as a source of noise in the analysis.

To generate random mixture fractions for the sampled cells, we sampled from Dirichlet distributions. Dirichlet distributions are the conjugate priors of multinomial distributions. Thus, a Dirichlet distribution is a distribution, with vector valued parameter $\gamma$, of probabilities for the multinomial distribution – in other words of mixture fractions for copy numbers obtained by sampling. For each region, each cell $i$ of the two dominant cells was assigned $\gamma_i = 10$, each cell $j$ of the 23 other selected cells from the same region was assigned $\gamma_j = 0.1$, and each cell $\ell$ of the other 50 cells was assigned $\gamma_\ell = 0.01$.

Fig. S1 shows the overall experimental design, including different regions from which single cells are collected and DNA sequenced. We extracted the copy number for each genomic locus from the SCS results to compose the cell matrix, as shown in the heatmap in the Fig. S1.

More formally, we chose $k$ total single cell samples, $\hat{k}$ dominant cell samples per region, and $\hat{n}$ simulated bulk samples per region. Here, $k = 75$, $\hat{k} = 2$ and $\hat{n}$ is 1, 2 or 3. We drew $\hat{n}$ column vectors of mixture fractions for each region from a Dirichlet distribution using the parameters $\gamma$, assigned as described above, for a total of $n = 3\hat{n}$ columns of mixture fractions. We use these $n$ columns to form the $k \times n$ matrix $\boldsymbol{F}^{(sel)}$. The $m = 9934$ and $k = 75$ columns of copy numbers

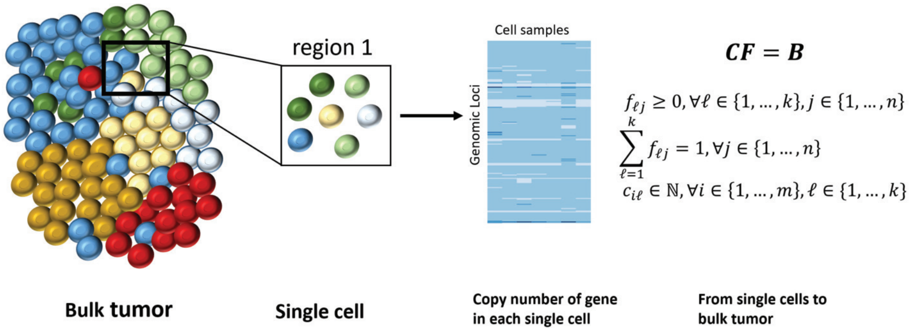


**Fig. S1.** The hierarchical structure of the bulk tumor and SCS data used for validation, and a basic example to compose a bulk tumor from single-cell data.

from the single cell data are used to form the $m \times k$ matrix $\boldsymbol{C}^{(sel)}$, and the $m \times n$ simulated matrix $\boldsymbol{B}$ is the product $\boldsymbol{B} = \boldsymbol{C}^{(sel)}\boldsymbol{F}^{(sel)}$. The entries of $\boldsymbol{B}$ are real values, not rounded to integers.

We also selected a number $k_{observed} = 3\hat{k}$ of single cells to form the matrix $\boldsymbol{C}^{(observed)}$ for use in some algorithms. The number of single cells selected varied, but in each case, the same number of cells was selected from each region, and the cells were selected from those that were not chosen to be among the 75 used to simulate the bulk tumor.

Our goal is to recover the $k' = 3\hat{k}$ (here $k' = 6$) dominant cells, representing the major clones from the simulated bulk samples. With this notation, the problem can be formally stated as follows:

Input: A matrix of bulk tumor $\boldsymbol{B}_{m\times n}$ and a matrix of observed single cell sequencing samples $\boldsymbol{C}^{(observed)}_{m\times k_{observed}}$.
Output: A set of inferred fundamental cell components $\boldsymbol{C}^{(inferred)}_{m\times k'}$ and corresponding set of mixture fraction $\boldsymbol{F}^{(inferred)}_{k'\times n}$.

#### A.5 Assessing Solution Quality

Given $\boldsymbol{B}_{m\times n}$, $\boldsymbol{C}^{(observed)}_{m\times k_{observed}}$. and $k'$, we would like to find $\boldsymbol{C}^{(inferred)}_{m\times k'}$ and $\boldsymbol{F}^{(inferred)}_{k'\times n}$ such that:

$$\boldsymbol{B}_{m\times n} \approx \boldsymbol{C}^{(inferred)}_{m\times k'}\boldsymbol{F}^{(inferred)}_{k'\times n.}$$

We repeated the above sampling, simulating, and deconvolving procedure for $N = 40$ experiments to assess the performance. Fig. 2 summarizes the design by which we simulated bulk tumor from the existing SCS data and how we integrated the SCS data into bulk tumor deconvolution.

We used two ways to estimate the performance. We calculated an accuracy, measured as the fraction of genomic positions (among the 40 replicates) with a correctly inferred copy number relative to the true cell components. We also measured the accuracy of inferred cell components and mixture fractions by the root mean squared deviation (RMSD) between true and inferred data.

For each genomic position in each inferred clone, if the copy number is equal to the copy number in the true clone, we consider the value to be accurately inferred. Then, we calculate the copy number error for each genomic position as follows:

$$\boldsymbol{C}^{(acc)}_{m\times k'} = \boldsymbol{C}^{(inferred)}_{m\times k'} - \boldsymbol{C}^{(true)}_{m\times k'.}$$

We define the accuracy to be the fraction of 0's in $\boldsymbol{C}^{(acc)}$.

Following [31], we also calculate the RMSD of $\boldsymbol{C}^{(inferred)}_{m\times k'}$ and $\boldsymbol{C}^{(true)}_{m\times k'}$, specified as the root mean square distance over all entries of all cell components between the two matrices:

$$\sqrt{\sum_{i=1}^{m}\sum_{j=1}^{k'}\left(c_{ij}^{(true)} - c_{ij}^{(inferred)}\right)^2 / mk'}.$$

Similarly, we can measure the RMSD between $\boldsymbol{F}^{(inferred)}_{k'\times n}$ and $\boldsymbol{F}^{(true)}_{k'\times n}$ over all the loci and mixture fractions:

$$\sqrt{\sum_{i=1}^{k'}\sum_{j=1}^{n}\left(f_{ij}^{(true)} - f_{ij}^{(inferred)}\right)^2 / k'n}.$$

Following the same idea, when we assess the RMSD between inferred results and true data in each cell component, we calculate the RMSD in the pair-wise columns of the matrices:

$$\sqrt{\sum_{i=1}^{m}\left(c_{ij}^{(true)} - c_{ij}^{(inferred)}\right)^2 / m}, \text{ for } \forall j \in \{1, ..., k'\}.$$

$$\sqrt{\sum_{j=1}^{n}\left(f_{ij}^{(true)} - f_{ij}^{(inferred)}\right)^2 / n}, \text{ for } \forall i \in \{1, ..., k'\}.$$

In the inferred trees of thirteen nodes each (six inferred, six observed, and a diploid root), we tested for clustering by tumor region as follows. Let the two-element sets $\{x_1, x_2\}, \{y_1, y_2\}, \{z_1, z_2\}$ be the nodes representing the two observed cells selected from the first region, second region, and third region respectively. Considering the inferred tree as an undirected tree, we can define unique undirected paths between $x_1$ and $x_2$, between $y_1$ and $y_2$, and between $z_1$ and $z_2$. A pair of nodes/cells representing one region is considered to "cluster together" relative to another region if the path between the two nodes from the region of interest does not pass through either of the two nodes from the other region.

##### A.6 Fully Simulated SCS Data

We conducted additional tests on fully simulated data in order to provide some test case for which the ground truth is known and for which data could be distributed without restriction. We modeled these fully simulated data on the real data to match the true number of regions $d$ ($d = 3$ in our case), number of cells $c_i$ ($i \in \{1, 2, 3\}$) per region, estimated rate $r_{ai}$ of copy number variation $a$ per region ($a \in \{0, 1, 2, ..., 10\}, i \in \{1, 2, 3\}$), and probability $p_{mi}$ that each genomic position that has a non-diploid copy number ($m \in \{1, 2, ..., 9934\}, i \in \{1, 2, 3\}$).

For each region $i$, we created a root with diploid copy number in all genomic positions. We then created a binary tree where each node represents one clone. We created copy number vectors for the nodes of the tree by adding mutations according to a Poisson distribution with empirical rate $r_{ai}$ and $p_{mi}$ to mutate the copy number in different genomic positions so that the overall copy number distribution would be similar that of the real SCS samples. We chose the depth $D$ of the tree to be the minimum value that allows the number of nodes to exceed the number of cells sampled in the real SCS data in each region (in our case, $D = 6$).

From the full tree, we constructed sub-trees starting from the root with $c_i$ nodes (excluding the root) by random walk with depth-first order and with the equal possibility to go right or left (Fig. S2). We collected the chosen nodes to establish an artificial single-cell data set encoded as a matrix whose rows are the genomic positions and columns are the simulated single-cell samples. We applied the methods described in Sec. A.4 on these simulated SCS samples to simulate bulk tumor samples, which we call the fully simulated tumor data.

We then further perturbed the artificial single-cell data to evaluate sensitivity of the methods to noise in CNA calling. We randomly choose a desired fraction (0, 0.2, or 0.4) of total genomic positions in each cell sample of $\boldsymbol{C}^{(observed)}$ to perturb, then sampled a set of uniformly random genomic loci without replacement and increased or decreased (if the copy number is greater than 1) the copy number by 1. This modification resulted in noisy versions of $\boldsymbol{C}^{(observed)}$. We then applied our methods and quality assessment as described in Sec. 3.1, Sec. 3.2 and Sec.A.5 to noiseless and noise-added variants of the fully simulated data.

#### B Supplementary Results

##### B.1 Inference Quality on Semi-Synthetic Bulk Data via Pure NMF

In this subsection, we describe further analysis and additional experimental results on the use of pure NMF or simple heuristic extensions thereof. We assessed the methods with a series of experiments on semi-synthetic data designed to assess the phylogeny-free and phylogeny-based methods in comparison to one another and to generic NMF as functions of various parameters of the simulated data and the algorithms. In general, NMF is used to decompose high-dimensional

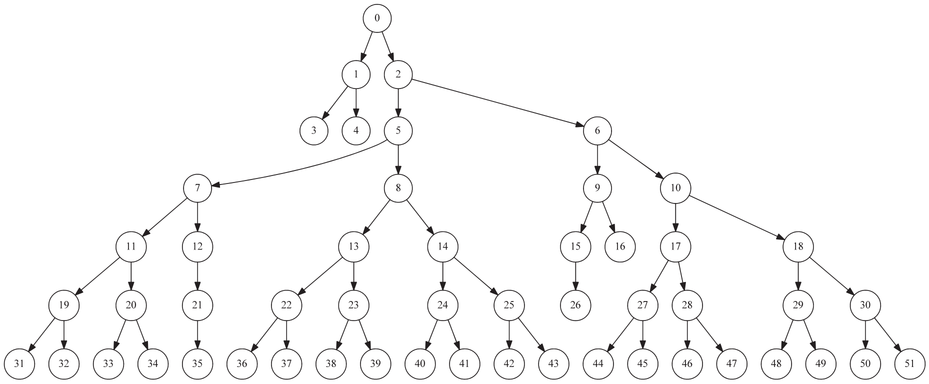


**Fig. S2.** A sub-tree chosen from the simulated completed binary tree of an example tumor region in the fully simulated data. The root was included in the tree structure for this visualization, but would be excluded when it was converted to a copy number matrix for use as input to our method.

data $(m \times n)$ to a low rank basis $(m \times k)$ and fractions $(k \times n)$, where we usually have $k << min(m, n)$. Therefore, we tested the pure NMF algorithm on different numbers of tumor samples to assess the data needs with respect to numbers of tumor samples. For these tests, we explored a wider range of sample numbers (3-99) than in the main paper to get a sense of whether performance improvement saturates for unrealistically large numbers of samples. As described in the preceding section, we assume that we have $k' = 6$ dominant cell components (derived from 2 out of random 25 cells selected in each region) as well as 23 rare components and assess our ability to infer the dominant components, treating the remainder as noise.

The copy number is predominantly diploid (2) across genomic positions for these data. We therefore trivially achieve relatively high accuracy either in each cell component or in all clones by guessing that all clones are purely diploid. As shown in the left plot in Fig. S3, the accuracy in some cell components can reach as high as 80%. The overall average accuracy resulting from setting all genetic positions to be diploid, indicated by the dashed line, means that over 70% of the entries are diploid. The right plot in Fig. S3 shows that the overall RMSD in copy numbers is also small for an all-diploid baseline solution. The variance means that in each clone, there are some loci of which the copy numbers are far away from diploid, as shown in Fig. 4 and Fig. 6. However, the all-diploid baseline is otherwise uninformative, since we are specifically interested in studying CNAs that change copy number for a subset of the genome. Nevertheless, we can use the accuracy of the all-diploid solution as a baseline against which to assess our algorithms.

We first applied NMF with random initialization to decompose the bulk tumor to infer the clonal cell components and the fraction matrix. We extended the test from small (3) to large (99) tumor samples with the regularization parameter $\alpha$ changing from 0 to 1 with the increment of 0.2. The $\alpha = 0$ results

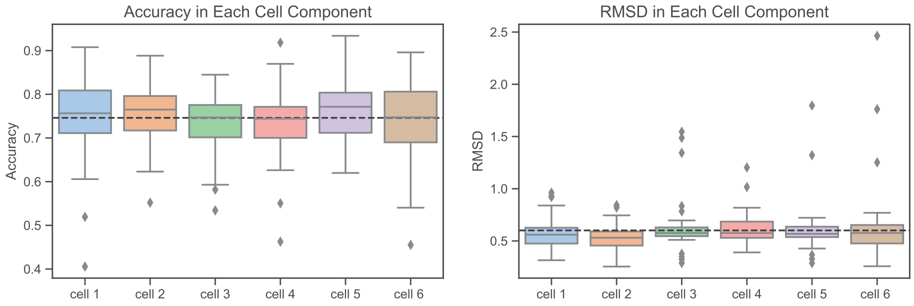


**Fig. S3.** Results of average and variance of accuracy (left) and RMSD (right) in copy numbers in each cell component. In this analysis, we compared a diploid cell matrix with all entries equal to 2 to the true cells in all the replicates to get the average and the variance of the all-diploid baseline solution. The results show that copy numbers are predominantly diploid in all clones, although with some variance. The black dashed line indicates the overall average value among all clones. Each colored box indicates one cell component. The Y-axis shows the accuracy or RMSD, respectively.

in the top three plots in Fig. S4 shows that the pure NMF method performs poorly in resolving the problem. For some $\alpha$, we can achieve a good estimation of accuracy and RMSD in copy numbers in smaller tumor samples (3, 6, 9) but the improvement is generally minimal for larger numbers of tumor samples (33, 99), although some specific $\alpha$ values can return good estimates. These results indicate that adding the penalty to the objective has the potential to lead to a good local optimization. We also note the performance of mixture fraction inference does not change much either across different numbers of tumor samples or among different $\alpha$.

The bottom three plots in Fig. S4 show results of applying the full phylogeny-free method, with real SCS data for initialization and objective penalty, to a larger range of sample sizes. The results of smaller tumor samples (3, 6, 9) have been shown in the main paper, but here we include the results from larger numbers of tumor samples (33, 99). As mentioned in the main paper, using real SCS data in initialization and the objective function together can lead to good estimates of copy numbers and mixture fractions of clones. Increasing the number of tumor samples substantially above the intrinsic dimension of the mixture does not appreciably improve the peak accuracy but does appear to make the method more robust to variation in $\alpha$.

#### B.2 Inference Quality on Semi-Synthetic Bulk Data via Phylogeny-Free Augmentation with SCS Data

Although we showed in the main manuscript that the phylogeny-free method can yield reasonably good average results (Fig. 3), we were further interested in how performance of the method might vary across cell components inferred. Thus, we calculated the accuracy and RMSD of inferred and true clones pairwise (Fig. S5)

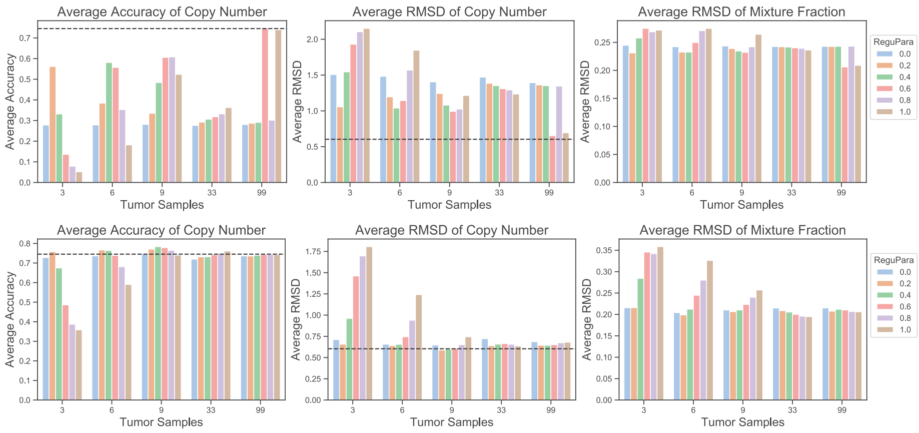


**Fig. S4.** NMF decomposition with $L_2$ penalty with random initialization (top) and with SCS initialization (bottom) for varying numbers of tumor samples. The figure shows the overall average accuracy of copy number (left), average RMSD in copy number (center), and average RMSD in mixture fraction (right) for deconvolution of varying numbers of tumor samples ($n$=3, 6, 9, 33, 99) for random initialization and initialization with true SCS data as a function of varying numbers of tumor samples and varying regularization parameter for $L_2$ penalty for deviation between true and inferred single cells. The black dashed lines in the left and center columns show accuracy and RMSD of copy number assignment for all-diploid inferences.

to see if the improvement is consistent in each clone. We also assessed this performance with varying regularization parameter $\alpha$ from 0 to 1 in increments of 0.2. We only show the results for smaller tumor samples (3, 6, 9) since previous results have shown no significant difference for larger numbers of tumor samples. The top plot of Fig. S5 shows that fine-tuned $\alpha$ can improve the accuracy of copy numbers in each clone while continuously increasing $\alpha$ would result in worse performance in each clone. Similar results can also be observed in RMSDs of copy numbers (center, Fig. S5) and RMSDs of mixture fraction (bottom, Fig. S5). These results indicate that the modified NMF method affects the performance for each individual clone rather than only having the effect on a subset of the clones.

#### B.3 Inference Quality on Semi-Synthetic Bulk Data via Phylogeny-Based Augmentation with SCS Data

This subsection elaborates on the results for the phylogeny-based method of Sec. 2.3, also specifically in evaluating accuracy and RMSD of cell components and mixture fractions in each cell component as functions of the regularization parameter $\beta$. Fig. S6 shows the assessment of performance in each cell component with varying regularization parameter $\beta$ from 0 to 1 in increments of 0.2. The top and center rows in Fig. S6 show that the accuracy and RMSDs in copy numbers were poor in each cell component when $\beta = 0$, but were substantially

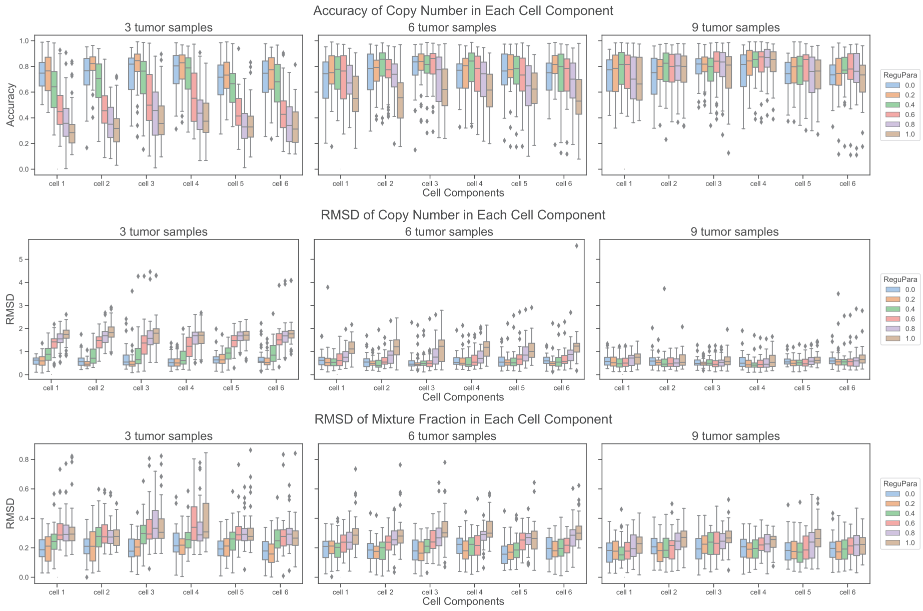


**Fig. S5.** Performance assessment by cell component for the phylogeny-free method. The results are from the same experiments as in Fig. 3 but analyzed in each cell component. The top row shows the accuracy of copy number in each component. The center row shows RMSDs of copy numbers in each cell component. The bottom row shows RMSDs of mixture fraction in each cell component. Columns (left to right) show results for varying numbers of tumor samples ($n = 3, 6, 9$).

improved by adding the phylogeny into the objective. Each cell component is fairly robust to $\beta$ variation and shows substantially better copy number accuracy and RMSD than is observed with pure NMF ($\beta = 0$). However, the RMSDs of mixture fractions (bottom row, Fig. S6) do not show appreciable improvements relative to pure NMF. Examination of the variances of the mixture fractions showed that adding the phylogeny increased the variances, which suggests that combining other objective function terms with the phylogeny might lead to better estimates of the fractions for each cell component.

#### B.4 Comparison of Phylogeny-Free and Phylogeny-Based Methods

We next compared the performance in each cell component of the two methods for three, six and nine tumor samples. We chose the best-performing regularization parameter, as indicated in Fig. 7, for each method in each tumor sample respectively. When comparing the results in each cell component, we found that the phylogeny-based method consistently gave better average accuracy and copy number RMSD for each cell component for three tumor samples (top and center row, Fig. S7). Also, the number of outliers from the phylogeny-based method

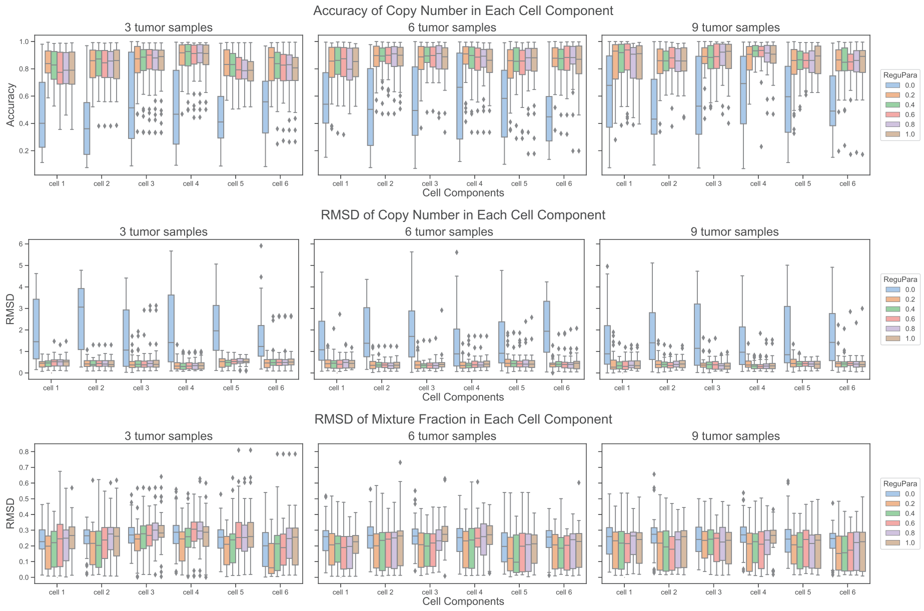


**Fig. S6.** Performance assessment by cell component for the phylogeny-based method. The results are from the same experiments as in Fig. 5 but analyzed in each clone. The top row shows the accuracy of the copy number predictions in each component. The center row shows RMSDs of copy number predictions in each cell component. The bottom row shows RMSDs of mixture fraction predictions in each cell component. Columns (left to right) show results for varying numbers of tumor samples ($n = 3, 6, 9$).

is generally smaller than that from phylogeny-free method in most of the cell components. In contrast, for mixture fraction comparison, the phylogeny-based method yields wider variances. This observation indicates the performance is less consistent in mixture fraction inference, but given the higher mean accuracy can be interpreted as an ability to infer much better mixture fractions in a subset of cases. (bottom row, Fig. S7).

#### B.5 Inference Quality on Fully Synthetic Data

In this section, we present the results of both phylogeny-free and phylogeny-based methods on fully synthetic tumor samples (Sec. A.6) following procedures similar to those of Sec. 3.1 and 3.2. We varied numbers of tumor samples (3, 6, 9) for these tests and regularization parameters ($\alpha = 0.0, 0.2, 0.4, 0.8$ for the phylogeny-free method and $\beta = 0.0, 0.2, 0.4, 0.8$ for the phylogeny-based method). For the phylogeny-free method (top row, Fig. S8), the results are qualitatively similar to those on semi-simulated data (Fig. 3) across the range of chosen regularization parameters. The phylogeny-free method shows notable improvement over pure NMF for at least some regularization parameters, with the range of effective regularization parameters expanding with increasing numbers of tumor

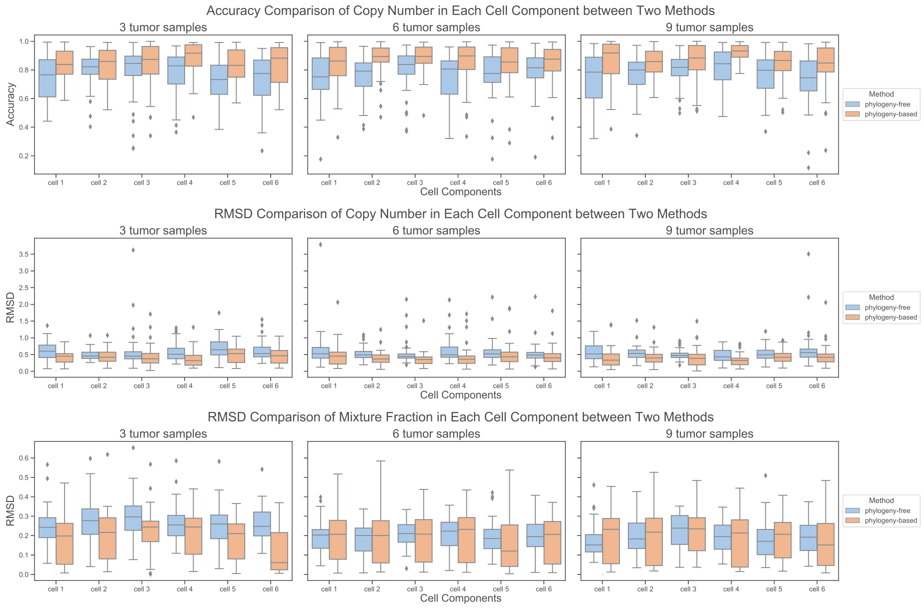


**Fig. S7.** Comparison in each cell component between phylogeny-free and phylogeny-based methods. The figure shows the comparison between the two methods for incorporating SCS data, using optimized regularization parameters for both as indicated in Fig. 7. Results are from the same experiment as in Fig. 7 but analyzed in each component. The top row shows the comparison of accuracy of copy number predictions in each component. The center row shows the comparison of RMSDs of copy number predictions in each cell component. The bottom row shows the comparison of RMSDs of inferred mixture fraction in each cell component. Columns (left to right) show results for varying numbers of tumor samples ($n = 3, 6, 9$).

samples. As with the semi-simulated data, the results of the phylogeny-based method (bottom row, Fig. S8) for $\beta > 0$ are substantially better than those for the phylogeny-free method. Further, they show limited much less sensitivity to sample size and to regularization parameter across the range of non-zero values considered. The phylogeny-based results on fully simulated data are somewhat better than those seen on semi-simulated data, suggesting that the more complicated error profiles of real SCS data do present some challenge to the methods.

Fig. S9 shows sensitivity of the methods to noise in $\boldsymbol{C}^{(reference)}$ at noise levels 0, 0.2, and 0.4. We conducted these tests for a single regularization value of 0.2 for $\alpha$ or $\beta$, chosen because it yielded relatively insensitive performance to sample size in the noise-free tests. For the phylogeny-free method (top row, Fig. S9), the plot shows that accuracy by each measure generally degrades with increasing noise, with copy number inference generally more sensitive to noise than mixture fraction inference. Robustness to noise by each measure increases noticeably with larger sample sizes. For the phylogeny-based method (bottom row, Fig. S9), the plot shows that accuracy is stable with increasing noise for

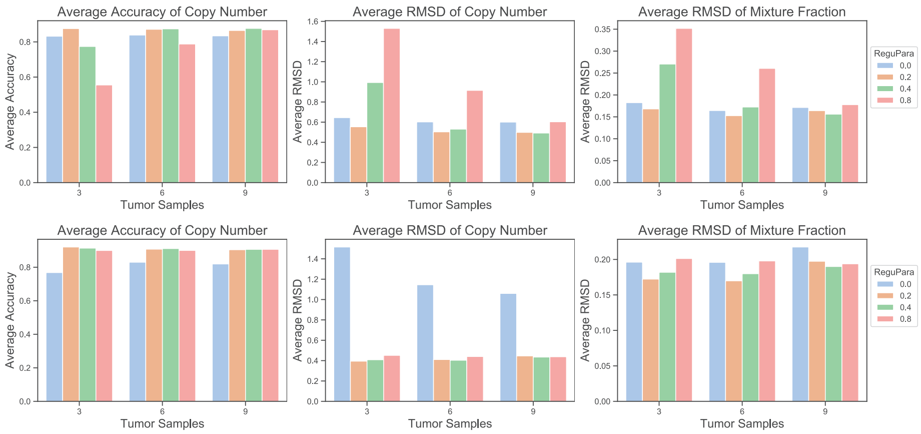


**Fig. S8.** Average accuracy and RMSD for our two methods as functions of tumor samples and regularization parameter on fully simulated tumor data. The top row shows the results for the phylogeny-free method, while bottom row shows the results for the phylogeny-based method. The left column shows the average accuracy of inferred copy numbers, the center column average RMSD between inferred and true copy numbers, and the right column average RMSD between the inferred and true mixture fractions. Bar plots show performance with regularization parameters 0.0, 0.2, 0.4 and 0.8. The X-axis shows the number of tumor samples and the Y-axis the average accuracy or RMSD.

all cases. RMSD results show some variability parameter to parameter, but no evident trend with increasing noise. These results show that the phylogeny-free method is robust to noise for sufficiently large sample size, while the phylogeny-based method is robust to noise even for small sample sizes.

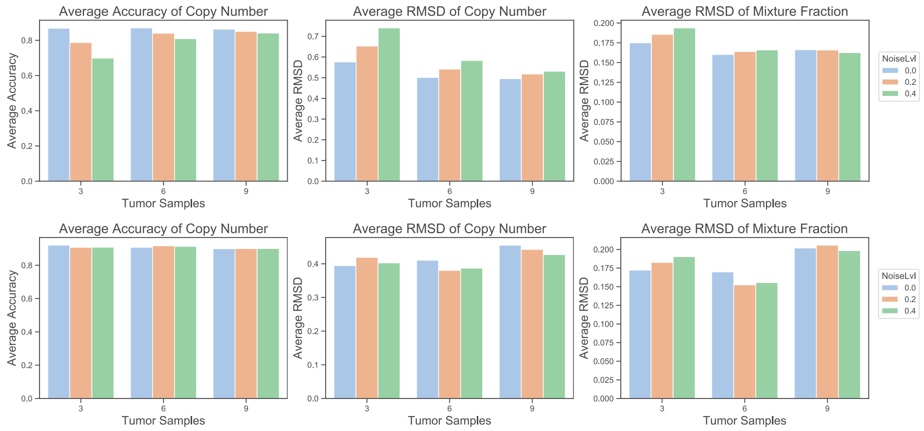


**Fig. S9.** Average accuracy and RMSD of sensitivity analysis on phylogeny-free (top) and phylogeny-based methods (bottom) as functions of noise level and number of tumor samples. The top row shows the results for phylogeny-free method, and the bottom row shows the results for phylogeny-based method. The left column shows the average accuracy of inferred copy numbers, the center column average RMSD between inferred and true copy numbers, and the right column average RMSD between the inferred and true mixture fractions. Bar plots show performance at different noise levels 0.0, 0.2, and 0.4. The X-axis shows the number of tumor samples and the Y-axis the average accuracy or RMSD.
